# SLIDE-VIP: a comprehensive, cell line- and patient-based framework for synthetic lethality prediction in DNA damage repair, chromatin remodeling and cell cycle

**DOI:** 10.1101/2022.07.07.499118

**Authors:** Magda Markowska, Magdalena A Budzinska, Anna Coenen-Stass, Senbai Kang, Ewa Kizling, Krzysztof Kolmus, Krzysztof Koras, Eike Staub, Ewa Szczurek

## Abstract

Discovering synthetic lethal (SL) gene partners of cancer genes is an important step in developing cancer therapies. However, identification of SL interactions is challenging, due to a large number of possible gene pairs, inherent noise and confounding factors in the observed signal. To discover robust SL interactions, we devised SLIDE-VIP, a novel framework combining eight statistical tests, including a new patient data-based test iSurvLRT. SLIDE-VIP leverages multi-omics data from four different sources: gene inactivation cell line screens, cancer patient data, drug screens and gene pathways. We applied SLIDE-VIP to discover SL interactions between genes involved in DNA damage repair, chromatin remodeling and cell cycle, and their potentially druggable partners. The top 883 ranking SL candidates had strong evidence in cell line and patient data, 250-fold reducing the initial space of 200K pairs. Drug screen and pathway tests provided additional corroboration and insights into these interactions. We rediscovered well-known SL pairs such as RB1 and E2F3 or PRKDC and ATM, and in addition, proposed strong novel SL candidates such as PTEN and PIK3CB. In summary, SLIDE-VIP opens the door to the discovery of SL interactions with clinical potential. All analysis and visualizations are available via the online SLIDE-VIP WebApp.

## Introduction

Synthetic lethality (SL) is classically defined as an interaction of two genes, where the co-inactivation of both genes results in cellular death, while inactivation of each individual gene results in a viable phenotype [1]. Discovering synthetic lethal gene pairs is an important step in developing new targeted cancer therapies [1, 2]. The mechanism that leads to a therapeutic window for SL-based therapies in cancer is that inactivation of one gene in the SL pair already occurs via the endogenous mutation in the tumor cells, and not in the normal cells throughout the rest of the body. Thus, applying a drug that targets the SL partner of that gene is expected to selectively kill cancer cells, leaving the normal cells viable.

The first approved targeted cancer therapy inhibiting the DNA damage response harnesses the SL interaction between *BRCA1* or *BRCA2* (genes often mutated in breast and ovarian cancer) and *PARP1/2* genes [1, 3]. Therapies using *PARP* inhibitors exhibit enhanced efficacy in patients with pre-existing *BRCA* deficiencies. Other ongoing clinical trials screen drugs targeting genes involved in DNA damage response (DDR), such as *ATR* inhibitors, for patients with *ATM*-mutated cancers [2, 4–6]. Further efforts to discover novel SL interactions are needed to propose new candidate pairs and develop new SL-based biomarker-driven cancer therapies.

With the assumption of around 20 thousand genes in the human genome, the space of potential SL partners consists of around 200 million potential pairs. Thus, experimental discovery of new SL partners without previous computational screening is infeasible. On the other hand, statistical analysis of all potential gene pairs can return many false positives. In our work, we aim at finding potential SL interactions that can be used for cancer therapy. Thus we focus on genes that take part in molecular processes important for cancer development and progression (further referred to as *focus genes*) or can be considered as therapeutic targets (*druggable genes*). The focus gene set comprises genes related to DDR, chromatin remodeling and cell cycle. The DDR pathway includes genes that are responsible for recognizing and repairing DNA damage in the cell [7], such as known tumor suppressors *TP53*, *ATM*, *ATR*, *BRCA1* and *BRCA2*. Genes from chromatin remodeling pathway, such as *ARID1A*, *ARID1B*, *SMARCA2* and *SMARCA4* from the SWI/SNF complex, encode proteins forming molecular machinery that alters chromatin structure and thus takes part in regulating gene expression as well as DNA repair [8]. Mutations in cell cycle genes (for example *TP53*, *RB1*, *CDKs* or transcription factors from *E2F* family) can deregulate the cell cycle, which is an indication of four cancer hallmarks: Self-Sufficiency in Growth Signals, Insensitivity to Antigrowth Signals, Limitless Replicative Potential and Genome Instability [9].

The computational methods for detecting SL gene pairs can be classified according to the input data source they utilize. The first type of methods, such as SLIdR [10], Synlet [11], or DepMap [12] exploit high-throughput gene dependence screens performed on cell lines, generated by large experimental efforts such as project DRIVE [13] or DepMap [14]. The second group of methods analyzes cancer patients survival and tumor genomic data (gene expression or somatic alterations [15]). Pipelines such as MiSL [16], DAISY [17], DiscorverSL [18] or ISLE [19] combine several statistical methods to analyze different types of pan-cancer tumor and cancer cell line data. Finally, many methods do not directly use experimental data, but mine databases containing biological networks [20], gene ontologies, protein-protein interactions or known SL interactions and employ machine learning techniques to detect patterns similar to those present for the known SL gene pairs [21, 22].

Integrating multiple statistical tests performed on several independent data sources is expected to provide better statistical power and increase the robustness of SL discovery from the imperfect and noisy biological data. In this study, we devised SLIDE-VIP (Synthetic Lethality Integrated Discovery Engine - Verified In Patients), a framework for SL interaction discovery, combining eight statistical tests, including a novel, patient data-based test iSurvLRT. Our framework leverages multi-omics data from four different sources: gene inactivation (knockout and knockdown) cell line screens (two tests on two different datasets), cancer patient data (four tests), drug screens (one test on two datasets) and genomic pathways (one test; Figure 1). We carefully assess the quality of the outcomes for each of them, rank the results with combined and adjusted p-values and choose potential SL pairs that are best supported by the most reliable results. Additionally, we offer an easily available SLIDE-VIP WebApp with pre-computed results and visualizations of eight SL tests for over 220 thousand gene pairs. By closely examining the known and newly discovered SL pairs we show that our comprehensive framework has the potential to discover new biomarkers or targets for cancer therapy.

**Figure 1:**
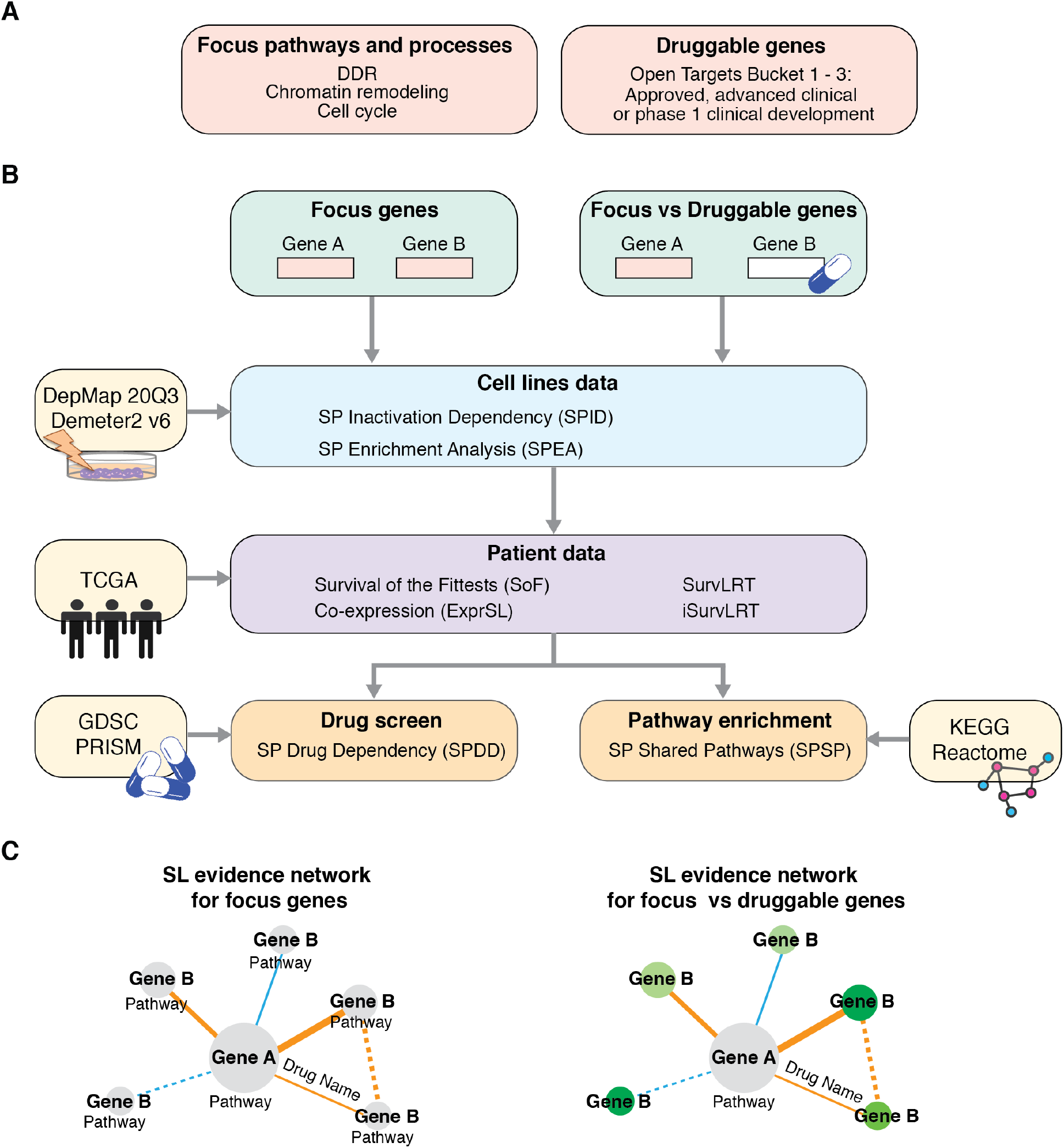
Synthetic lethal partners identification workflow used in SLIDE-VIP. The focus genes were defined based on their involvement in one of the focus pathways and processes, and druggable genes were selected from the Open Target database (Bucket 1-3). **A** The two input sets of gene pairs (focus genes and focus genes vs druggable genes) were then tested in cell line data (DepMap20Q3 and Demeter2 v6), followed by the statistical analysis of patient data (TCGA), drug screen validation (GDSC and PRISM) and pathway enrichment (KEGG and Reactome). **B** The results were utilized to create SL evidence networks summarizing the evidence from all tests. **C** The color of the node illustrates the drug development stage (dark green approved, light green clinical and yellow preclinical development stage). GDSC, the Genomics of Drug Sensitivity in Cancer; KEGG, Kyoto Encyclopedia of Genes and Genomes; PRISM, Profiling Relative Inhibition Simultaneously in Mixtures; SL synthetic lethality; SP synthetic partner; TCGA, The Cancer Genome Atlas.

## Methods

### Definitions

#### LoF alteration

For both cell line and patient data, we considered loss of function (LoF) alterations that result in producing a non-functional protein. LoF alterations were defined based on the Ensembl [23] classification of the variants, and only variants with a predicted high impact effect were utilized. This included the following alterations: splice site, nonsense and damaging missense mutations, frameshift insertion/deletion, start codon insertion/deletion/single nucleotide polymorphism, stop codon insertion/deletion. To assess the functional effect of the missense variants and to define damaging ones, we used ANNOVAR [24]. Five variant effect prediction methods included in ANNOVAR were used to increase the accuracy of the prediction and decrease the false-positive rate: PolyPhen-2 [25], SIFT [26], PROVEAN [27], CADD [28] and DANN [29]. A missense mutation was classified as damaging if at least three out of these five methods predicted a damaging effect on the protein function according to the damaging score cut-off established by each method.

#### Focus genes

Focus genes were defined as genes from DDR, chromatin remodeling and cell cycle pathways. Sets of genes involved in these pathways were collected based on published literature [30–35] and curated databases (total 1,241 genes; Supplementary Table 1; Figure 1A).

#### Druggable genes

Druggable genes were defined as genes targeted by drugs in an advanced stage of development. Specifically, 953 druggable genes were selected from the Open Targets Platform database ([36], release 19.06), based on the stage of development of the drug that targets the selected gene, including genes in the approved, advanced clinical and phase 1 clinical development stage (Bucket 1 - 3, Supplementary Table 2).

### Data source and pre-processing

#### Cell line data

The dataset of genome-scale CRISPR knockout screens from the project Achilles (https://depmap.org/portal/achilles/) for 789 cell lines was obtained from the Cancer Dependency Portal (DepMap 20Q3 release [37], https://depmap.org/portal/download). 787 of these cell lines had genomic information and thus were utilized in the analysis. Additionally, large-scale RNAi screening datasets including the Broad Institute Project Achilles, Novartis Project DRIVE [13] and the Marcotte et al. [38] breast cell lines, with the genetic dependencies estimated using the DEMETER2 model [39], were downloaded from the same source, the DepMap portal (712 cell lines). The genomic characterization data of the cell lines was generated by the Cancer Cell Line Encyclopedia (CCLE) and obtained from DepMap 20Q3 (787 and 670 cell lines for the Achilles and DEMETER2 projects, respectively). The cell line alteration data was then organized into a cell line per gene binary matrix with the value 1 if the cell line acquired at least one LoF alteration in the gene, and 0 otherwise.

#### Patient data

The somatic SNV (single nucleotide variants) alterations, mRNA gene expression and clinical data for cancer patients were downloaded for The Cancer Genome Atlas (TCGA) Pan-Cancer via the University of California Santa Cruz Xena data portal [40]. Raw counts from TCGA expression data were log2-scaled and quantile normalized to mitigate batch effects between different datasets (e.g. sequencing lab, time, etc.). Quantile normalization was performed using the “preprocessCore” R package [41]. The patient data was organized into a patient per gene binary matrix with the value 1 if the patient acquired at least one LoF alteration in the gene, and 0 otherwise.

#### Drug screen data

The drug screening data for the drug sensitivity analysis was collected from the Genomics of Drug Sensitivity in Cancer database (GDSC; https://www.cancerrxgene.org/downloads) [42–44], and the Profiling Relative Inhibition Simultaneously in Mixtures database (PRISM [37]) from the DepMap data portal. Specifically, we exploited both the two available GDSC experimental setups (GDSC1 and GDSC2) and the secondary PRISM Repurposing 19Q4 screen. GDSC datasets were curated using the Open Targets database [36] to unify the gene and protein information and assign gene targets to each drug. The screens included 518 unique drugs (367 in GDSC1 and 198 in GDSC2, with some overlap) targeting genes B in 988 unique cell lines in the GDSC database (987 cell lines in GDSC1 and 809 cell lines in GDSC2, with some overlap) and 1210 drugs targeting genes B in 481 cell lines in PRISM database.

#### Pathway data

The gene pathways were collected from the curated gene sets: Kyoto Encyclopedia of Genes and Genomes (KEGG [45–47]), Pathway Interaction Database (PID [48]) and Reactome [49], available as part of the c2 collection in MSigDB [50]. Together, we evaluated 1951 pathways: 186 in KEGG, 196 in PID and 1569 in REACTOME.

### Synthetic lethality tests

We used eight tests on four data sources (cell line, patient, drug and pathway data) to collect evidence of synthetic lethality: two types of statistical tests on cell line data, four tests on patient data, two tests on drug screens and one pathway enrichment test.

#### Synthetic Partner Inactivation Dependency (SPID)

SPID test is an application of a one-sided Wilcoxon rank-sum test to determine whether cell lines with LoF alteration in gene A have significantly lower gene B dependency scores than the cell lines without gene A LoF alteration, i.e. whether they are more sensitive to gene B inactivation. In addition, we calculate the *positive dependency percentage* which is the fraction of such cell lines out of the total number of cell lines, which have a positive estimated dependency score as a result of gene B deactivation. A positive dependency score implies that the cells grow and divide more efficiently when gene B is deactivated. Thus, a high positive dependency percentage implies that a given pair should not be considered SL.

##### Test application criteria

To perform SPID test for a given gene pair, gene A must have LoF alterations in at least 20 cell lines and gene B must be a knock-out target in a given cell line experiment.

##### SL evidence criteria

Gene pairs with the combined and adjusted p-value less than 0.05, where the combination is across two datasets (Achilles and DEMETER2) and two tests (SPID and SPEA).

#### Synthetic Partner Enrichment Analysis (SPEA)

SPEA test is a novel modification of a widely known and used Gene Set Enrichment Analysis (GSEA). GSEA identifies sets of genes that are over- or under-represented in lists ranked by gene expression [50]. Specifically, given a list of genes ranked by correlation of their expression with a certain phenotype of interest and a certain set of genes, GSEA uses a permutation-based test and a Kolmogorow-Smirnoff statistic to compute the significance of enrichment of the gene set at the top of the ranked gene list.

Instead of gene expression, SPEA ranks cell lines according to their gene dependency score of gene B (instead of differences in expression between two samples typically used in GSEA) so that the cell lines most sensitive to gene B silencing are situated at the top of the list. We are then interested whether the subset of such cell lines that carry gene A LoF alteration, is enriched at the top of that ranked list of cell lines. We thus calculate an enrichment score (ES) by walking down the list, increasing a running-sum statistic when we encounter a cell line from the subset, and decreasing it when we encounter cell lines without gene A LoF alteration. The ES is the maximum deviation from zero encountered in the random walk and corresponds to a weighted Kolmogorov–Smirnov-like statistic [50]. The ES indicates the direction of enrichment - a positive score means that cell lines with gene A LoF alteration are concentrated at the beginning of the ranking i.e. are more sensitive to gene B knockdown. See Supplementary Text for a formal description of the ES calculation. We estimate the statistical significance (nominal p-value) of the ES by comparing it with the set of scores ES_rand_, computed with randomly assigned gene A LoF alteration status and for a reordered cell line list. We perform this permutation step 200 times, recompute ESrand of the gene set for the permuted data and compile all results to generate a null distribution for the ES. The empirical, nominal p-value of the observed ES is then calculated relative to this null distribution.

##### Test application criteria

To perform SPEA test for a given gene pair, gene A must have LoF alterations in at least 20 cell lines and gene B must be a knock-out target in a given cell line experiment.

##### SL evidence criteria

Gene pairs with the combined and adjusted p-value less than 0.05, where the combination is across two datasets (Achilles and DEMETER2) and two tests (SPID and SPEA).

#### Gene co-expression (ExprSL)

ExprSL test is computed as the pairwise gene expression correlation and its significance is evaluated using a two-sided t-test for Spearman correlation. ExprSL is based on two assumptions. First is that SL gene pairs are involved in closely related biological processes and thus are more likely to be significantly positively correlated. Second is that the SL partner genes may compensate for each other and thus can be significantly negatively correlated [51].

##### Test application criteria

To perform ExprSL test for a given gene pair, both gene A and gene B’s expression must be measured.

##### SL evidence criteria

Gene pairs with the absolute Spearman correlation coefficient higher than 0.4 and adjusted p-value less than 0.05 are considered significantly correlated.

#### Survival of the Fittest (SoF)

SoF test assumes that cells with reduced expression of both genes in a given SL pair will not survive in a tumor cell population [51]. Thus, intuitively, SoF test assesses whether tumors with LoF alteration of gene A compensate for this loss by an increase of gene B expression. Specifically, SoF uses a one-sided Wilcoxon rank-sum test to examine whether gene B has a significantly higher expression in samples with LoF alteration in gene A compared to the rest of the samples.

##### Test application criteria

To perform SoF test for a given gene pair, gene A must have LoF alterations in at least 20 patients and gene B’s expression must be measured.

##### SL evidence criteria

Gene pairs with SoF test adjusted p-value less than 0.05 are considered significant according to this test.

#### SurvLRT

We use the survival likelihood ratio test (SurvLRT) [15] to estimate the tumor fitness with a given genotype g from survival data of patients. Here, the genotype *g* = (*g_A_*, *g_B_*) is defined by alterations in gene A and gene B. Specifically, for a given patient, *g_A_* = 0 if gene A is not altered in that patient’s tumor, and *g_A_* = 1 otherwise, and similarly for *g_B_*. In the original approach [15], the alteration could be of any type. Here, it is strictly confined to the definition of LoF alteration. SurvLRT assumes a survival model of tumor fitness, stating that a decrease in tumor fitness due to LoF alteration in gene A and gene B is exhibited by a proportional increase of survival of the patients. Thus, the survival of patients with LoF alterations in both SL genes should be longer than expected from the survival of patients without LoF alteration in those genes or with only one gene altered.

Consider a reference survival function *S*(*t*), estimated based on a cohort of patients who did not die of cancer as by [15]. Denote the fitness of a tumor with genotype *g* by Δ*_g_* and denote the log fitness as *δ_g_* = log(Δ*_g_*).

We assume that the survival of patients whose tumor carries genotype g is given by *S*(*t*)^Δ_*g*_^. In the case when there is no epistatic relation between genes A and B, we expect

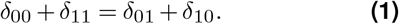

In the case when gene A and gene B are in any epistatic relation (positive or negative), however, we expect that

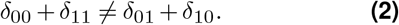

Finally, for A and B being synthetic lethal partners, we expect that

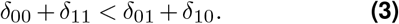

SurvLRT is a likelihood ratio test, which is based on analytical estimates of the parameters 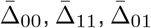, and 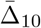, (and, correspondingly 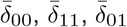, and 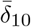,) and verifies the null hypothesis given by Eq. S (1) against an alternative hypothesis defined by inequality Eq. S (2). To decide if the detected interaction is synthetic lethal, we compute the effect size 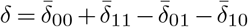 and check if the constraint *δ* < 0 holds. If it holds, we set an “SL flag” to 1 and report the pair as synthetic lethal with the associated p-value from the likelihood ratio test. Otherwise, we set the “SL flag” to −1. For each investigated pair, we in addition report the log fitness of the double LoF alteration genotype that would be expected in the case of no epistatic interaction, i.e., in the case when Eq. S (1) would hold, denoted 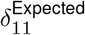.

Importantly, not all such SL interactions are clinically relevant. Inequality Eq. S (3) can also hold when *δ*_00_ < *δ*_11_. In such a case, the fitness of the genotype with double LoF alterations Δ_11_ is still unexpectedly high, given the single LoF alterations. In particular, it is smaller than the fitness of the double genotype expected in the case of no epistatic interaction, 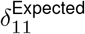. Such an interaction, however, is not clinically relevant: turning the synthetic lethal partner off in addition to the first gene in the pair using treatment would cause the patients to survive worse than patients without the inactivation of either of the genes. Thus, we in addition set a clinically relevant (“CL”) flag, which is 1 if *δ*_00_ > *δ*_11_ and −1 otherwise.

##### Test application criteria

To perform SurvLRT for a given gene pair, gene A must have LoF alterations in at least 20 patients. Additionally, at least 5 patients must have LoF alteration and be classified as deceased for each genotype.

##### SL evidence criteria

Gene pairs with SurvLRT test adjusted p-value less than 0.05, both SL flag and CL flag equal to one are considered significant evidence for clinically relevant SL according to this test.

#### iSurvLRT

Here, we propose a novel test, called iterative SurvLRT (iSurvLRT), which is an extension to SurvLRT. SurvLRT test [15] has limited applicability, as it can only be used to test such gene pairs where both genes carry LoF alterations in a sufficient number of tumor samples. Instead, for a given gene A, which is often altered in cancer, it is desirable to find a partner B that itself does not necessarily acquire alterations in tumors.

The solution is iSurvLRT that bases on LoF alterations in gene A and expression of gene B. Specifically, LoF alteration status of gene A in a given patient is defined as *g_A_* = 0 if A is not altered in the patient and *g_A_* = 1 if gene A is altered. The expression status of gene B in the same patient is defined based on the fact whether the expression of gene B, denoted *e_B_*, is low in that patient or not, i.e. *g_B_*(*t*) = 0 if *e_B_* >= *t* and *g_B_*(*t*) = 1 if *e_B_* < *t* for a given threshold *t*. To define the threshold *t*, we consider a grid of possible thresholds given by the quantiles of the empirical distribution of expression of gene B. Specifically, the grid of thresholds is defined by *t* ∈ (*q*(0.05), *q*(0.1),…, *q*(0.5)), where *q*(*α*) is the *α*-th quantile of the distribution of *e_B_*. We next iterate over the grid of thresholds and define the genotypes *g*(*t*) = (*g_A_*, *g_B_*(*t*) for each patient based on the current threshold *t* in the iteration. For each iteration, we compute the p-value in SurvLRT test for the genotype *g*(*t*) obtained for the current threshold. Finally, we return the lowest p-value across all iterations, the threshold used in that iteration and the obtained SL and CL flags.

##### Test application criteria

To perform iSurvLRT for a given gene pair, gene A must have LoF alterations in at least 20 patients and gene B’s expression must be measured. At least 5 patients must have LoF alteration and be classified as deceased for each genotype.

##### SL evidence criteria

Gene pairs with iSurvLRT test adjusted p-value less than 0.05, both SL flag and CL flag equal to one are considered significant evidence for clinically relevant SL according to this test.

#### Synthetic Partner Drug Dependency (SPDD)

SPDD test assesses the susceptibility of cancer cell lines with gene A LoF alteration to drugs targeting gene B. To this end, cell lines are grouped to either wild type (WT) or gene A LoF. The potential drug sensitivity is assessed using the drug targets from GDSC and PRISM data and a one-sided Wilcoxon test on the LNIC50 values (natural logarithm of the fitted half maximal inhibitory concentration for GDSC datasets) and median-collapsed log fold change profile corresponding to the WT and LoF group (cell lines without or with LoF alteration in gene A, respectively). This test is performed separately for each drug and the drug with the best result (the lowest p-value) is reported.

##### Test application criteria

To perform SPDD test for a given gene pair, gene A must have LoF alterations in at least 20 cell lines and gene B must be a known drug target.

##### SL evidence criteria

Gene pairs with SPDD test p-value less than 0.05 are considered significant evidence for SL according to this test.

#### Synthetic Partners Shared Pathways (SPSP)

SPSP assesses enrichment in common pathways using a part of c2 collection from MSigDB, consisting of pathways from KEGG, PID and Reactome databases. Specifically, a hypergeometric test is used to detect pairs that share a significantly large number of pathways, given the number of pathways each of the genes is involved in.

##### Test application criteria

To perform SPSP test for a given gene pair, both gene A and gene B must be present in at least one of the considered pathways.

##### SL evidence criteria

Gene pairs with SPSP test adjusted p-value less than 0.05 are considered significant evidence for SL according to this test.

### slideCell and slidePat - cell line and patient tests R implementation

SL tests on cell line data were implemented in an R package called slideCell. Given dependency scores and alteration information for cancer cell lines, as well as a list of gene pairs, slideCell can be used to generate statistics, p-values and plots for SPID and SPEA tests. The source code for slideCell package is freely available at [slideCell](https://github.com/szczurek-lab/slideCell). SoF, ExprSL, SurvLRT and iSurvLRT tests on patient data were implemented in an R package slidePat, which takes as input gene alteration and expression, as well as patient survival data. The source code for slidePat package is freely available in [slidePat](https://github.com/szczurek-lab/slidePat).

### SLIDE-VIP WebApp

The SLIDE-VIP WebApp is an online application for the visualization of test results for 224,169 potential SL gene pairs from this publication. The application has been developed in RStudio [52], version 1.2.5033, using the *Shiny* package, version 1.5.0 and R [53] version 4.0.5. The application is freely available online at slidevip.app.

## Results

### SLIDE-VIP – a workflow for identifying synthetic lethal partners of genes involved in DNA damage response, chromatin remodeling and cell cycle from integrated cell line, patient, pathway and drug screen data

We devised a uniquely comprehensive workflow for identifying clinically relevant synthetic lethal interactions (Figure 1). We performed two series of synthetic lethality tests (Figure 1B). In the first series, we tested pairs of focus genes (genes involved in DNA damage response, chromatin remodeling and cell cycle; Methods and Supplementary Table 1). In the second series, we tested pairs of focus genes versus druggable genes (focus genes versus genes that have known targeting drugs in advanced stages of development; Methods and Supplementary Table 2). For each gene pair, gene A was chosen from the focus genes and assessed if at least 20 LoF alterations were present in the dataset (see Methods for a definition of LoF alteration). The second gene in the pair, referred to as gene B, was either a focus or a druggable gene, and it was considered a candidate for a synthetic lethal partner of gene A.

First, a total of 224,169 gene pairs in both series (126,175 focus gene pairs and 97,994 focus vs druggable gene pairs) were assessed for SL interactions using SPID and SPEA tests based on the Achilles and DEMETER2 knock-out screen data for cell lines (see Methods). We only analyzed pairs that were possible to test on both datasets. Next, we combined the obtained four p-values using Fisher method [54] and then performed Benjamini-Hochberg multiple testing correction [55] for the resulting combined p-value for all 224,169 gene pairs (together for the focus and focus vs druggable genes’ pairs). With that procedure we obtained 24,193 pairs (14,556 focus, 9,637 focus vs druggable) with combined p-value < 0.05, and 1,833 pairs (1,267 focus and 566 focus vs druggable) with combined and corrected p-value < 0.05. We consider the 1,833 pairs to be the top cell line-based SL candidates. Next, we screened the 224,169 gene pairs for confirmation in the patient data, drug screens and pathway enrichment. First, four patient tests were performed using 7,256 tumor samples in TCGA. Since each of the tests has different application criteria (see Methods) we were able to apply each of them to a different number of pairs (see table 1). It is worth noting that introducing a novel iSurvLRT test allowed us to use patient survival data for a much higher number of pairs than with the existing SurvLRT test. We then performed Benjamini-Hochberg multiple test correction for each of the four patient tests for the all tested pairs (together for focus and focus vs druggable genes’ pairs, see table 1). Finally, we checked which of the top 1,833 cell line-based pairs showed evidence of SL for at least one of the patient tests (see Methods for SL evidence criteria). In this manner, we obtained 883 clinically relevant SL gene pairs (683 focus and 200 focus vs druggable) that we consider the top SL candidates with strong evidence in both cell line and patient data.

**Table 1:** Summary of the performed SL tests. The number of gene pairs tested using each test differs because the tests have different application criteria and are performed on different datasets. Additional test criteria for gene pairs: *SL flag and CL flag in SurvLRT and iSurvLRT tests equal to 1; **absolute correlation for ExprSL test higher than 0.4; ***using the adjusted p-value.

| SL Test | Dataset | No. of gene pairs tested | No of pairs with corrected p-value < 0.05 and additional SL criteria | No of performed tests among the top 1,833 cell line based | No of pairs with additional validation among the top 1,833 cell line based |
| --- | --- | --- | --- | --- | --- |
| SPID and SPEA | Achilles and DEMETER2 | 224,169 | 1,833 | - | - |
| SoF | TCGA | 200,405 | 68,571 | 1,714 | 647 |
| SurvLRT | TCGA | 23,478 | 2,434* | 376 | 50* |
| iSurvLRT | TCGA | 191,801 | 10,464* | 1,668 | 89* |
| ExprSL | TCGA | 197,058 | 13,831** | 1,702 | 204** |
| At least one patient test | TCGA | 200,405 | 86,589 | 1,714 | 883 |
| SPDD | PRISM | 62,026 | 12,293 (1***) | 536 | 98 (0***) |
| SPDD | GDSC | 26,354 | 5,833 (220***) | 351 | 115 (6***) |
| SPSP | KEGG, PID and REACTOME | 178,256 | 12,488 | 1,606 | 212 |

We checked for additional evidence for the top SL pairs in public cell line drug screening data (Methods). A relatively small number of pairs (62,026 pairs for PRISM dataset and 26,354 for GDSC dataset) included a gene B that was also targeted in publicly available drug screens (see table 1). After performing SPDD test on PRISM and GDSC dataset, we applied the Benjamini-Hochberg multiple test correction to the obtained SPDD p-values separately for each of the two datasets. The p-value histograms for both datasets (Supplementary Figure 1) show a strong skew of the unadjusted p-value distribution towards 0, indicating that for multiple tested pairs there is a signal for their SL interaction in that data. Accordingly, before the multiple testing correction, there were 5,833 pairs in GDSC dataset and 12,293 pairs in PRISM dataset (see table 1) with p-values less than 0.05. However, after the Benjamini Hochberg correction, there were only 220 pairs in GDSC dataset and one pair in PRISM dataset with the adjusted p-value less than 0.05. This resulted from the fact that, in contrast to the previous tests, the range of the original p-values does not include extremely low values. Since we consider this test as an additional validation rather than a means for identifying new pairs, we further considered the unadjusted p-values for that test. With that, we acquired 115 pairs among the top 1,833 cell lines based on GDSC dataset and 98 pairs for PRISM dataset (see table 1). Those pairs are especially interesting because drugs targeting gene B demonstrate potential for drug repurposing and targeted cancer therapy for patients with LoF alteration in gene A.

Finally, to gain additional insight into the biological function of the identified pairs, we evaluated 1,951 pathways from the c2 collection in MSigDB [50] database to identify significant common enrichment of potential SL partners in curated gene sets. We performed SPSP test for 178,256 gene pairs and then performed Benjamini-Hochberg multiple test correction. Among the top 1,833 cell line based pairs, 212 SL gene pairs shared a highly significant number of pathways, which suggests that their SL interaction can be explained by their shared functionalities. The common pathways for these pairs may be important for conveying the SL interactions, i.e., it is likely that the pair is SL because the pathways may operate when only one of the genes is turned off, but they fail to function when both are lost.

To rank the top 883 clinically relevant SL gene pairs, we used the corrected combined p-value for the cell line-based tests. We did not include the other tests in the combined p-value for the ranking because they could not be performed for all of the pairs and each of them had a different sample size, resulting in varying statistical power. All the results for the top pairs, sorted according to the final ranking, are presented in Supplementary Table 3 (683 focus pairs; referred to as focus ranking) and Supplementary Table 4 (200 focus vs druggable pairs; referred to as focus vs druggable ranking). Combined with plots illustrating all the performed tests (easily available through our webtool SLIDE-VIP WebApp) they are a rich source of knowledge about relevant SL interactions among genes from DDR, chromatin remodeling and cell cycle pathways and potentially druggable genes.

### Previously known SL pairs for focus genes

Several synthetic lethal interactions between the focus genes found by using SLIDE-VIP have already been identified previously, thus confirming the validity of our approach. Among the top 683 focus pairs, 30 are featured in the SynLethDBv2.0 database [56]. This is a very significant enrichment (p-value 7.5 × 10^−63^; hyper-geometric test with the assumption of 200 mln possible gene pairs and 37 thousand being featured in SynLethDBv2.0). Specifically, the four top pairs are already confirmed SL gene pairs: *TP53* and *CHEK2*, *RB1* and *E2F2*, *RB1* and *SKP2*, *TP53* and *PLK1*.

For DDR genes, the top identified focus pairs include known SL gene pairs such as: *TP53* and *CHEK2* (first in the focus ranking [57, 58]), *PRKDC* and *ATM* (15th in the focus ranking [59]) and *TP53* and *IGF2* (44th in the focus ranking [60, 61]. All these three pairs have very strong confirmation in cell line tests and are confirmed either by SoF or SurvLRT patient data test. For the cell cycle genes, we rediscovered known SL interactions between *RB1* and *E2F3* (second in the focus ranking [62–64]) and *RB1* and *SKP2* (third in the focus ranking [63, 65, 66]). Those two pairs have full confirmation in 4 cell line data tests and are also confirmed by SoF patient data test. Among the top ranking pairs, there was also one already known pair *TP53* and *PLK1* (fourth in the focus ranking [4, 67]) that has very good confirmation in DEMETER2 cell line data tests and also SoF patient test.

Previous work identified a set of 31 gene pairs, consisting of paralogs that share common functionalities and thus display SL interactions [22], as well as SL pairs in SWI/SNF chromatin remodeling complex deficient cancers [35]. Out of these 31 pairs, seven were among the 224,169 gene pairs tested using our framework. The remaining ones were excluded either because they did not belong to the focus or druggable sets, or did not pass the testing criteria. To enrich the body of knowledge of these seven pairs, we investigated in detail the type of evidence for their SL interactions that could be found using SLIDE-VIP. Three out of those seven were among the top 1833 cell line based pairs, six were confirmed by at least one test on patient data, three found additional confirmation using drug screens and four using pathway-based tests (table 2). In the SWI/SNF complex, the cell line tests confirmed the SL interaction for known paralog pairs *ARID1A* and *ARID1B* [13, 68, 69] and *SMARCA4* and *SMARCA2* [35, 70]. *ARID1A* and *ARID1B* synthetic lethal interaction was also observed in patients tests. *SMARCA4* and *AURKA* synthetic lethal interaction was confirmed both in cell line and patient data.

**Table 2:** Paralogs and known SL pairs in SWI/SNF. Confirmation in each test is in accordance with the SL evidence criteria described in Methods. 1 confirmed; 0 not confirmed; NA test could not be performed.

| Gene A | Gene B | Source ref | Among the top 1,833 cell line based | TCGA confirmation: SoF / SurvLRT / iSurvLRT / ExprSL | Drug SPDD confirmation: PRISM / GDSC | Pathway confirmation: SPSP | SynLethDBv2.0 score |
| --- | --- | --- | --- | --- | --- | --- | --- |
| ARID1A | ARID1B | [22, 35] | 1 | 1 / 0 / 0 / 1 | NA / NA | 1 | 0.2 |
| SMARCA4 | SMARCA2 | [35] | 1 | 0 / 0 / 0 / 0 | NA / 1 | 1 | 0.4 |
| CREBBP | EP300 | [22] | 0 | 0 / 0 / 0 / 1 | 0 / NA | 1 | 0.3 |
| ARID1A | ATR | [35] | 0 | 0 / 1 / 0 / 0 | 0 / 1 | 0 | 0.2 |
| SMARCA4 | AURKA | [35] | 1 | 1 / NA / 0 / 0 | 1 / 1 | 0 | 0 |
| ARID1A | EZH2 | [35] | 0 | 1 / 0 / 0 / 0 | 0 / 0 | 1 | 0.2 |
| ARID1A | PARP1 | [35] | 0 | 1 / 0 / 0 / 0 | 0 / 0 | 0 | 0 |

### Previously known SL pairs for focus vs druggable genes

In contrast to the top 683 focus gene pairs, the top 200 pairs of the focus vs druggable genes are not necessarily expected to be previously studied and known (Supplementary Table 4). The druggable partners may be involved in any type of other processes than the focus DDR, chromatin remodeling or cell cycle pathways. Thus, the focus vs druggable test series is more exploratory and intended to provide candidates for drug repurposing. For the top 200 focus vs druggable pairs, four were also found in SynLethDBv2.0 database [56]. This is still a significant enrichment (p-value 5.3 × 10^−10^) measured with a hypergeometric test with the assumption of 200 mln possible gene pairs and 37 thousand being synthetic lethal (the number of records in SynLethDBv2.0). Out of those, the best ranked are *TP53* and *MAPKAPK5* (ninth in the focus vs druggable ranking) and *TP53* and *ITGB1* (49th in the focus vs druggable ranking) with additional confirmation in SoF patient data test.

### SLIDE-VIP confirmed synthetic lethality interaction between *PRKDC* and *ATM*

We next selected one pair of particular interest from the top candidates for the focus SL gene pairs, namely the *PRKDC* and *ATM* pair (Figure 2). This pair has confirmed SL potential according to the SynLethDBv2.0 database [56]. The waterfall plot of *ATM* dependency in cell line knock-out screens shows enrichment of negative dependency scores in cell lines with *PRKDC* LoF (red bars in Figure 2A). For that pair, SPID test for the Achilles dataset identified that dependency of cell lines on *ATM* with *PRKDC* LoF alteration is significantly higher compared to cell lines without *PRKDC* LoF alteration (p-value 9.77 × 10^−6^; Figure 2B first plot). SPEA confirmed that the cell lines with *PRKDC* LoF alteration are significantly enriched at the top of the list of cell lines as ordered by their sensitivity to CRISPR-mediated knock-out *ATM* knock-out (p-value 0.005; Figure 2B first plot). Importantly, the same tests on the independent DEMETER2 dataset also confirmed the interaction (p-values 0.0009 for SPID and 0.005 for SPEA; Figure 2B and Figure 2C second plots). Using SurvLRT, we showed that patients with simultaneous LoF alterations of *PRKDC* and *ATM* survive significantly better than expected given the survival of patients with individual LoF alterations and without LoF alterations of those genes (p-value 0.004; Figure 2D). The evidence for synthetic lethal interaction between *PRKDC* and *ATM* was also found in drug data, where the sensitivity of cell lines with PRKDC LoF alteration to the drug KU-60019, known to target *ATM*, was significantly larger than of cell lines without *PRKDC* inactivation (p-value 0.043; Figure 2E). This gives additional drug-repurposing insight, that the drug KU-60019, originally developed as a drug targeting *ATM*, could be used to target cancers with *PRKDC* LoF alteration. Additional confirmation of the interaction between the genes was found in SPSP test, which shows that the genes share a significantly high number of pathways (*PRKDC* is part of 14 pathways, *ATM* is part of 51, and they have 5 common pathways in common; p-value 0.0004; Figure 2F).

**Figure 2:**
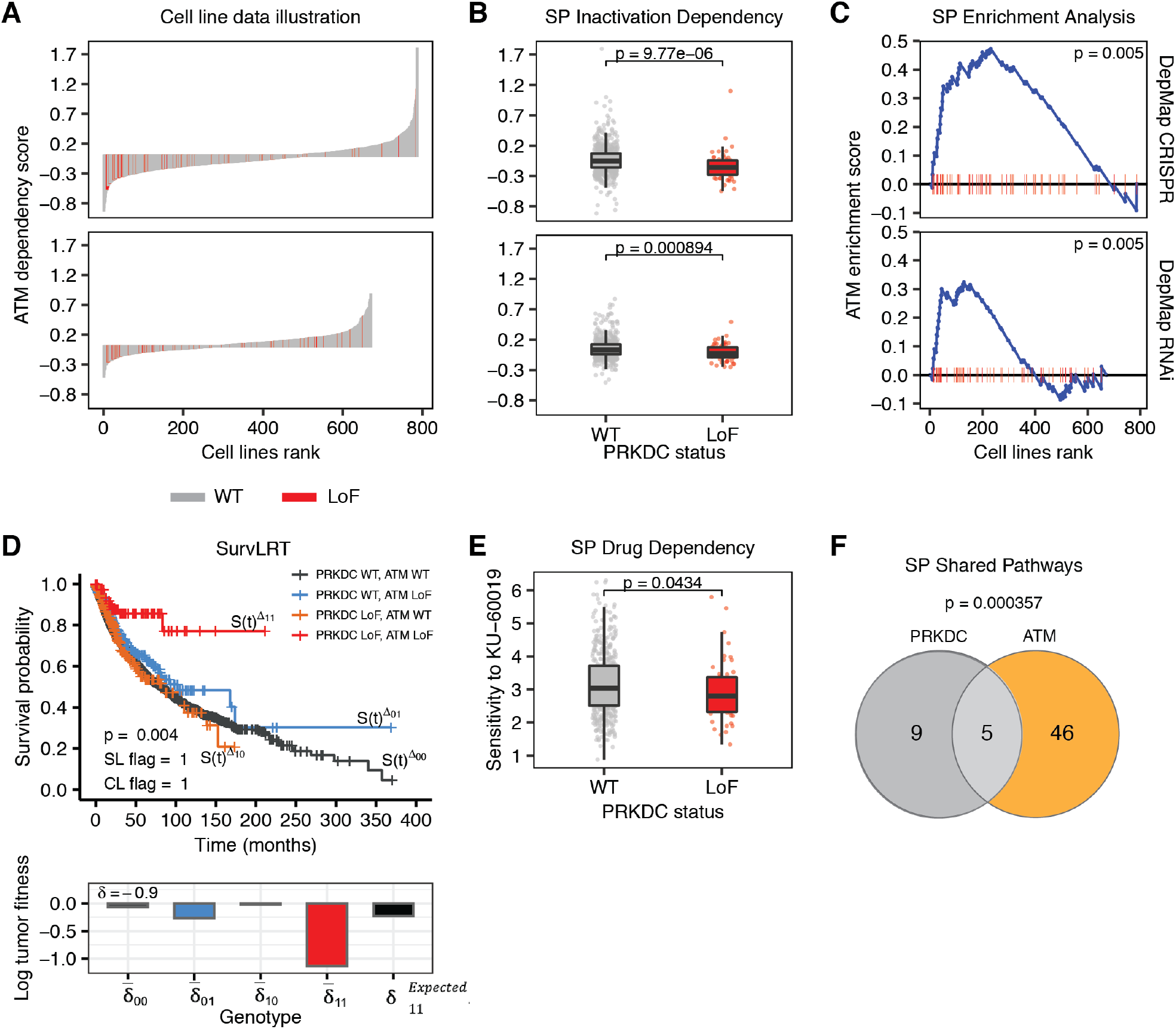
Evidence found in cell line, patient, drug and pathway data for the *PRKDC* and *ATM* SL interaction. Plots illustrating the results of tests that confirmed the SL interaction between the pair. **A** Waterfall plots for Achilles (first row) and DEMETER2 (second row) datasets, illustrating the results of *ATM* knock-out experiments in cancer cell lines (sorted according to the effect size, cell lines having *PRKDC* LoF marked in red; x axis) with knockout effect measured by dependency score (negative effect means that cells proliferate worse after the knock-out, positive - the cells proliferate better; y axis). **B** Boxplots illustrating SPID test results for Achilles (first row) and DEMETER2 (second row) datasets (y axis same as for panel A). **C** Plots illustrating SPEA test results for Achilles (first row) and DEMETER2 (second row) datasets; cell lines are sorted according to the dependency score (cell lines having *PRKDC* LoF marked in red; x axis); the blue line illustrates the calculations of *ATM* enrichment score (global maximum being the measured ES; y axis) that increases when encountering a cell line with *PRKDC* LoF, and decreases when encountering cell line with WT *PRKDC*. **D** The Kaplan-Meier survival plots illustrating SurvLRT test result with survival time in months (x axis) and calculated survival probability (y axis); the effect size for different *PRKDC* and *ATM* genotypes measured by log tumor fitness are shown below, 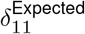 value representing the effect size expected without SL interaction for double *PRKDC* and *ATM* LoF genotype. **E** Boxplots illustrating SPDD test result for GDSC dataset; the cells sensitivity to drug targeting *PIK3CB* is measured by ln(IC50) (y axis). **F** Venn diagram illustrating SPSP test result with the number of common *PRKDC* and *ATM* pathways printed in the middle.

*PRKDC*, encoding the catalytic subunit of the DNA-dependent protein kinase DNA-PK, and *ATM* are both kinases capable of sensing DNA damage and activating downstream effector kinases to initiate the repair system. Loss of *ATM* has been tested in multiple clinical trials as a selection biomarker for drugs that target the DNA repair system such as Olaparib [71]. Pre-clinical research indicates that loss of *ATM* could sensitize cancer cells for DNA-PK inhibition [59, 72]. Given that DNA-PK was not profiled in the Depmap Achilles dataset, we could not see this previously described interaction, we rather see the reciprocal gene pair: loss of *PRKDC* as a sensitiser for *ATM* inhibition. Drugs targeting *ATM* have entered clinical development in recent years, so it would be of high interest to confirm this synthetic lethal interaction further.

### D*PTEN* and *ATM* pair is identified as a pair with high evidence for synthetic lethality for the focus vs druggable gene pairs

Another highly interesting candidate SL pair, *PTEN* and *PIK3CB*, was identified as second at the top focus vs druggable pairs ranking (Figure 3). The waterfall plot of *PTEN* dependency in cell line knock-out screens clearly shows enrichment of negative dependency scores in cell lines with *PIK3CB* LoF (red bars in Figure 3A). SPID test on both Achilles and DEMETER2 data indicates that cell lines with *PTEN* LoF alteration are significantly more sensitive to inactivation of *PIK3CB* (p-values 6.2 × 10^−6^ and 0.0004 respectively; Figure 3B). This is confirmed on the same datasets using SPEA test (both p-values equal to 0.005; Figure 3C). Finally, for patient data, iSurvLRT test found that patients with simultaneous *PTEN* LoF alteration and low expression of *PIK3CB* survive significantly better than expected from survival of patients with individual or without LoF alteration of *PTEN* or *PIK3CB* (p-value equal to 0.0004; Figure 3D), indicating high clinical relevance for that SL interaction. Drug data also shows evidence of synthetic lethal interaction between the genes, where the sensitivity of cell lines with *PTEN* LoF alteration to the drug AZD6482, known to target *PIK3CB*, was significantly larger than of cell lines without *PTEN* inactivation (p-value 7.4 × 10^−8^; Figure 3E). Drug AZD6482 is thus a strong candidate for SL-based repurposing, as this result suggests that AZD6482 should be further evaluated for the treatment of patients with *PTEN* LoF alteration. Additional confirmation of the SL interaction between the genes was found in SPSP test, which shows that the genes share a significantly high number of pathways (*PTEN* is present in 44 pathways, *PIK3CB* in 97, and they have 22 pathways in common; p-value 3.6 × 10^−15^; Figure 3F).

**Figure 3:**
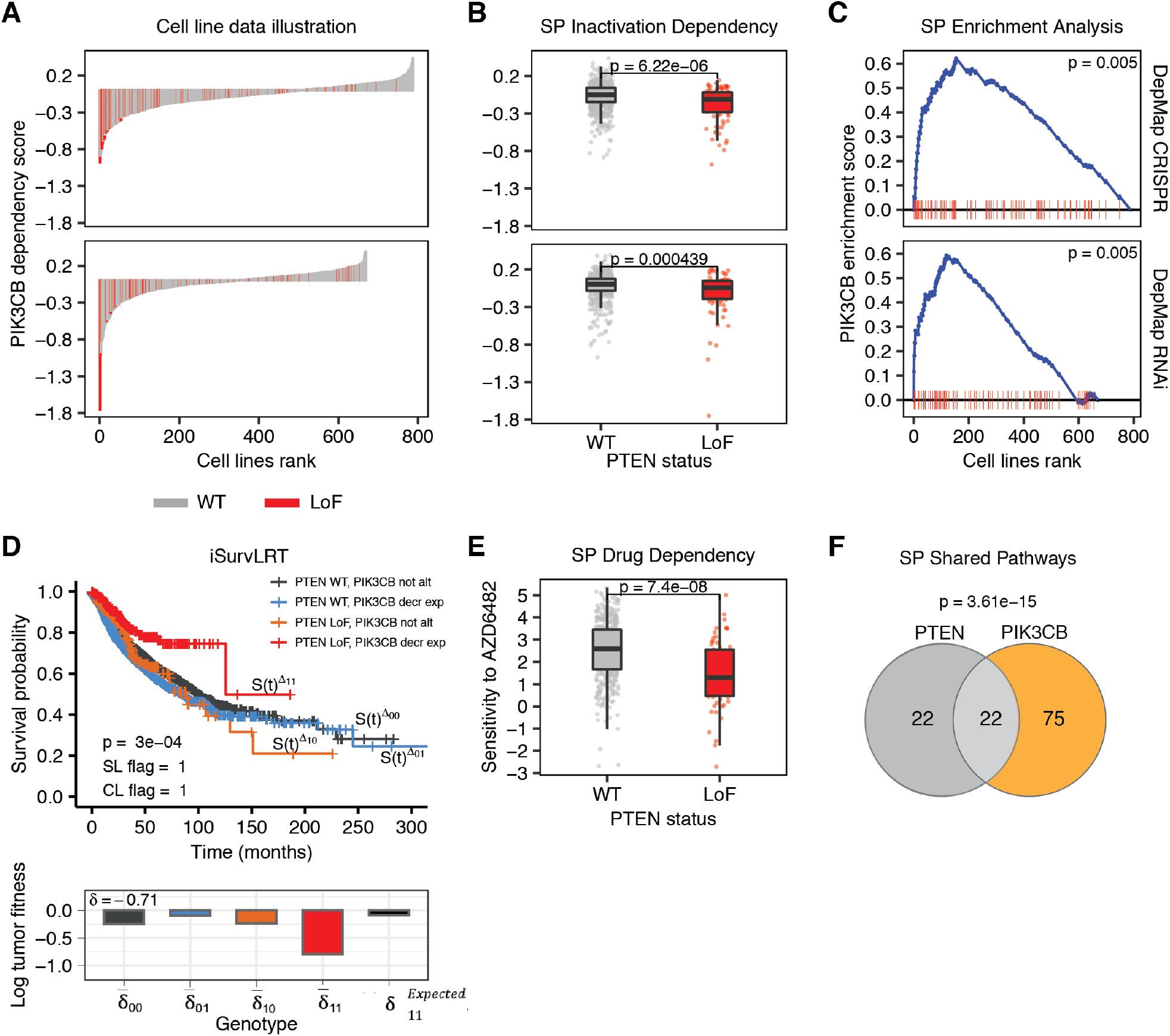
Evidence found in cell line, patient, drug and pathway data for the *PTEN* and *PIK3CB* SL interaction. Plots illustrating the results of tests that confirmed the SL interaction between the pair. **A** Waterfall plots for Achilles (first row) and DEMETER2 (second row) datasets, illustrating the results of *PIK3CB* knock-out experiments in cancer cell lines (sorted according to the effect size, cell lines having *PTEN* LoF marked in red; x axis) with knock-out effect measured by dependency score (negative effect means that cells proliferate worse after the knock-out, positive - the cells proliferate better; y axis). **B** Boxplots illustrating SPID test results for Achilles (first row) and DEMETER2 (second row) datasets (y axis same as for panel A). **C** Plots illustrating SPEA test results for Achilles (first row) and DEMETER2 (second row) datasets; cell lines are sorted according to the dependency score (cell lines having *PTEN* LoF marked in red; x axis); the blue line illustrates the calculations of *PIK3CB* enrichment score (global maximum being the measured ES; y axis) that increases when encountering a cell line with *PTEN* LoF, and decreases when encountering cell line with WT *PTEN*. **D** The Kaplan-Meier survival plots illustrating iSurvLRT test result with survival time in months (x axis) and calculated survival probability (y axis); the effect size for different *PTEN* genotype and *PIK3CB* expression level combinations measured by log tumor fitness are shown below, 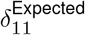 value representing the effect size expected without SL interaction for double *PTEN* and *PIK3CB* LoF genotype. **E** Boxplots illustrating SPDD test result for GDSC dataset; the cells sensitivity to drug targeting *PIK3CB* is measured by ln(IC50) (y axis). **F** Venn diagram illustrating SPSP test result with the number of common *PTEN* and *PIK3CB* pathways printed in the middle.

The *PIK3* pathway regulates a myriad of cellular processes including proliferation and survival by initiating phosphorylation signaling cascade [73]. The lipid phosphatase *PTEN* is a negative regulator of *PIK3* pathway activity. Consequently, deregulation of the *PIK3* pathway in tumors is frequently caused by inactivation or loss of *PTEN* [74, 75]. Interestingly, it has previously been observed that PTEN-deficient cancer cells require the lipid kinase activity of *PIK3CB* for both *PIK3* signaling and growth in vitro [76, 77], which gives additional support for that potential SL interaction.

### Synthetic lethality evidence network for focus genes

To visually inspect the evidence found for the most promising SL candidates and their interconnections, we created SL evidence networks for the top 50 pairs, both for the focus gene list (Figure 4) and for the focus vs druggable gene list (Figure 5). The evidence networks summarized all tests and data sources that confirmed the synthetic lethal interaction for each category of the pairs.

**Figure 4:**
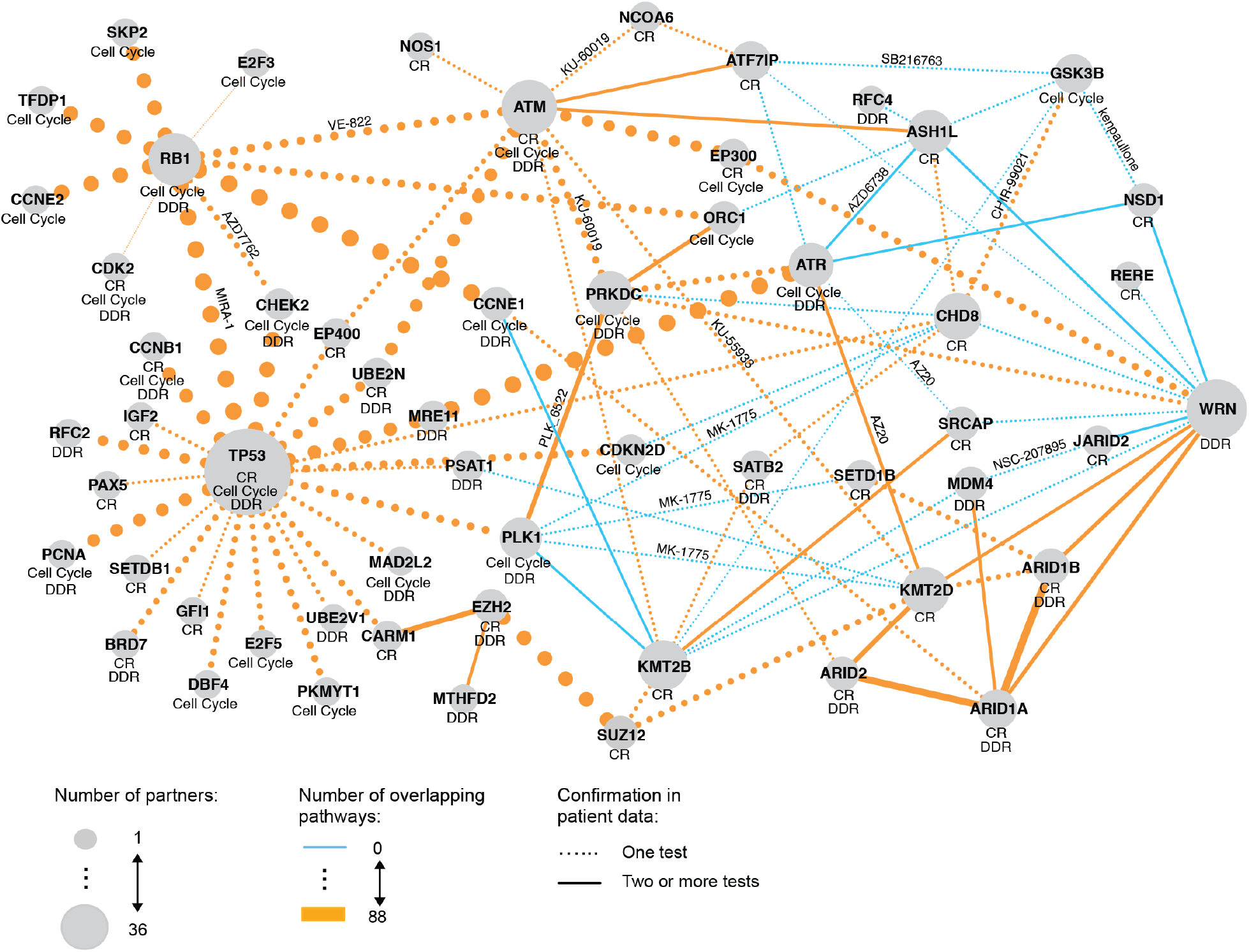
SL evidence network for focus genes. The SL evidence network summarizes the test results and data sources for the top 50 candidate pairs. Each node represents a gene (gene A or B) and each edge the number of overlapping pathways between two genes. The node size indicates the number of gene partners. The pathways (DDR, CR or cell cycle) in which a gene is involved are shown below the gene name. The drug name is shown next to the edge when the p-value of SPDD test was less than 5%. Confirmation from at least two tests from patient data is marked by a solid line. CR, chromatin remodeling; DDR, DNA damage response.

**Figure 5:**
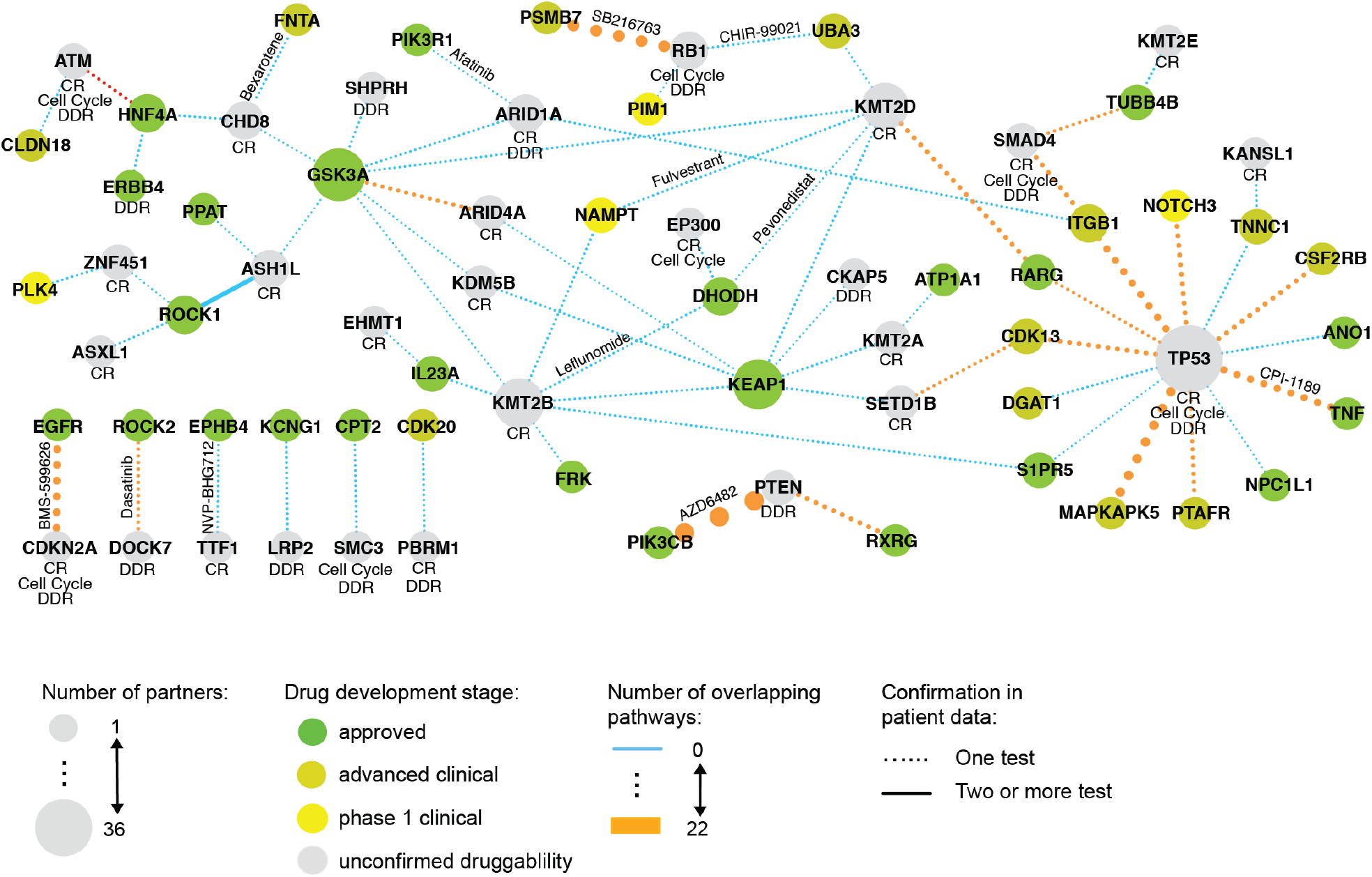
SL evidence network for focus vs druggable genes. The summary network for focus vs druggable genes for the top 50 candidate pairs. Each node represents a gene (gene A or B) and each edge width symbolizes the number of overlapping pathways between two genes. The color of the node represents the drug development stage and the node size indicates the number of gene partners. The drug name is shown next to the edge when the p-value of SPDD test was less than 5%. Confirmation from at least two tests from patient data is marked by a solid line. CR, chromatin remodeling; DDR, DNA damage response.

Interestingly, for the focus genes (Figure 4), a set of cell cycle genes forms a separated cluster, with *RB1* as a hub, involving genes such as *CDK2* or *E2F3* as partners. *TP53* emerged as the largest hub in the network, with a cluster of partners involved in chromatin remodeling (including *HDAC5*, *HDAC6* or *UBR2*). Around ATM, there was another, most densely connected cluster of genes involved in chromatin remodeling, which included *NOS1*, *KMT2D*, or *NCOA6*. Connected genes involved in the DDR include *ARID1B* and *ARID1A* (together with *ATR* via *KMT2D*). The network includes a link between *SMARCA2* and *SMARCA4*, paralogue genes known as synthetically lethal, involved in both chromatin remodeling and DDR.

Interestingly, the *WRN* gene formed an additional hub, with *PRKDC, ARID1A, ARID1B* and others as potential SL partners. Given *WRN* dependency is typically coupled to MSI status, these interactions could indicate additional vulnerabilities for MSI tumor cells.

### Synthetic lethality evidence network for focus vs druggable genes

Overall, compared to the SL evidence network for the focus genes, the SL evidence network for the focus vs druggable genes is less densely connected (Figure 5). However, there emerged several interesting hubs. First, again, *TP53*, a tumor suppressor involved in all three focus pathways, was identified as a SL partner of multiple druggable genes, 5 of which were already approved. Interestingly, two druggable genes, not involved in any of the focus pathways, emerged as hubs, which means they were identified as potential partners for many of the focus genes. In particular, *KEAP1* was found to be a candidate partner for five focus genes, four of which are involved in chromatin remodeling. *DHODH* was another popular partner, and was also identified in three pairs.

### The proposed workflow identifies novel potential SL pairs

We identified several novel interesting SL interactions relating to genes with roles in chromatin modification. For example, we observed that cell lines with LoF inactivation of the lysine methyltransferase 2D (*KMT2D/MLL4*) were more dependent on *ATM* than those with WT *KMT2D/MLL4*. Furthermore, evidence from SurvLRT, iSurvLRT and GDSC drug screening supported this potential SL interaction (Supplementary Table 3). Curiously, previous studies suggested that inactivation of *KMT2D* and also *KMT2C* increase sensitivity to Poly(ADP-ribose) polymerase inhibitors (PARPi) [78, 79], thus highlighting an important link between chromatin modification and targeting the DNA damage response. Moreover, *NCOA6* mutated cancer cell lines in GDSC screen exhibited a higher sensitivity to *ATM* inhibitor KU-60019 than those with WT *NCOA6*. Therefore this could be an interesting pair to investigate further experimentally.

We also observed that LoF alteration in the chromatin remodeling enzyme chromodomain helicase DNA-binding protein 8 (*CHD8*) sensitized cancer cell lines to inactivation of the exonuclease *MRE11*. Although this pair lacks confirmation from patient tests, it was supported by GDSC drug screen data where *CHD8* mutated cell lines were more susceptible to the MRN complex inhibitor Mirin.

## Discussion

Here, we introduced SLIDE-VIP – a novel comprehensive SL analysis framework, combining eight statistical tests and leveraging four data sources. It includes a novel, patient survival-based iSurvLRT test, which allows us to harness survival data for a higher number of potential pairs than using our previous SurvLRT test. Both iSurvLRT and SurvLRT inform about the clinical relevance of the tested SL interaction. Previous approaches for SL identification combining several data sources [18, 19] used less comprehensive workflows with a lower number of performed tests.

For a defined target space consisting of genes relating to DDR, chromatin remodeling and cell cycle, our framework indicated known and novel pairs with clinical relevance. The results obtained for the focus gene pairs are of different nature than those obtained for the focus vs druggable genes. The former comprehensively explored the interactions between the genes important for cancerogenesis. In this category we rediscovered many known SL gene pairs, at the same time revealing new ones that can be further looked into. The latter analysis – of the focus vs druggable gene pairs – was more exploratory and revealed a smaller number of cell line and patient verified pairs. Gene pairs from the second category may have a higher clinical potential with genes B being targeted with drugs that are in the advanced stage of development.

Conveniently, all our analysis results are publicly accessible via SLIDE-VIP WebApp. The plots illustrating the results are easily available for manual inspection by a specialist interested in developing new cancer therapies. To complete the picture, the app also shows the results of the additional tests on drug screens and pathway information.

Our framework must be viewed within the limitations of the available datasets. They are the results of either a biological experiment or data collected from the patients – both of those categories are prone to confounding factors and include noise in the data produced either on the experimental stage or during the bioinformatics preprocessing steps. They also have different sample sizes not only between the datasets but also inside them – for example, more cell lines and patients have LoF mutations in common cancer genes such as *TP53* or *RB1*, which can be part of the reason for their high position in the list ranked by combined p-values. Furthermore, the patient data is sparse, which we partly overcame by using four patient tests, including a novel one – iSurvLRT, that uses different types of input. Having higher sample sizes for each of the data types would allow us to apply SLIDE-VIP in a cancer indication specific manner. Cancer type is potentially an important predictive factor, especially for the survival data used in SurvLRT and iSurvLRT tests.

SLIDE-VIP uncovered gene pairs with excellent potential for developing new targeted cancer therapies. Our framework vastly reduces the space of candidate SL pairs and thereby enables focusing experimental verification on the most promising candidates. The pairs with additional verification in drug screens and pairs from the focus vs druggable genes are especially attractive for drug repurposing. In this way, SLIDE-VIP opens the door to the discovery of SL interactions with clinical potential.

## Data Availability

The sources of the data used in the framework are described in the Materials and Methods Section. The results for the top patient verified tests are summarized in the Supplementary Table 3 and 4. The results of all the performed tests are available at slidevip.app. The slideCell R package is available at [slideCell](https://github.com/szczurek-lab/slideCell). The slidePat R package is available at: [slidePat](https://github.com/szczurek-lab/slidePat).

## Author Contributions

MM, MB and ESz conceptualized the initial idea of the framework. MB and ESz developed the iSurvLRT methodology. MM performed the cell line tests and developed slideCell package, MB performed the patient data tests and implemented slidePat package, KKor and KKol performed the tests on drug data and EK performed the tests on pathway data. MM combined and ranked the results of all the tests. KKol implemented the SLIDE-VIP WebApp. MM, MB and EK prepared the figures. ACS and ESt provided biological interpretation of the results. MM, MB and ESz wrote the initial draft of the manuscript. All authors provided critical feedback; helped shape the research and analysis; edited, reviewed and approved the manuscript.

## Acknowledgements

The work by MM was done during Interdisciplinary doctorate studies using next generation sequencing in personalized medicine (at Postgraduate School of Molecular Medicine, Medical University of Warsaw).

