## Supplementary materials for "SLIDE-VIP: a comprehensive, cell line- and patient-based framework for synthetic lethality prediction in DNA damage repair, chromatin remodeling and cell cycle"

Supplementary information

Magda Markowska[1,2], Magdalena A Budzinska[1,3], Anna Coenen-Stass[4], Senbai Kang[1], Ewa Kizling[1], Krzysztof Kolmus[3], Krzysztof Koras[1], Eike Staub[4], and Ewa Szczurek[1,*]

[1] *Faculty of Mathematics, Informatics and Mechanics, University of Warsaw, Stefana Banacha 2, 02-097 Warsaw, Poland*

[2] *Postgraduate School of Molecular Medicine, Medical University of Warsaw, Zwirki i Wigury 61, 02-091 Warsaw, Poland*

[3] *Ardigen, Podole 76, 30-394 Kraków, Poland*

[4] *Translational Medicine, Oncology Bioinformatics, Merck Healthcare KGaA, Frankfurt Strasse 250, 64293 Darmstadt, Germany*

### Contents

1 SPEA (Synthetic Partner Enrichment Analysis) – formal description 3

Supplementary Figure 1 4

### 1 SPEA (Synthetic Partner Enrichment Analysis) – formal description

Let's denote a cell line as $cl$, number of cell lines as $n$, dependency score - $ds$, list of cell lines ranked by decreasing dependency score - $L$, a position on the list - $i$, set of cell lines with alteration - $C1 = c1_j : 1, 2, ..., n_a$, set of cell lines without alteration - $C0 = c0_j : 1, 2, ..., n - n_a$. Then the ES score is the (weighted) Kolmogorov-Smirnov (K-S) statistic defined as:

$$ES = \sup_{1<i<n} (F_i^{C1} - F_i^{C0})$$

The ES is the largest difference in $F$ which are the (weighted) empirical cumulative distribution functions:

$$F_i^{C1} = \frac{\sum_{t=1}^{i} |ds_t|^p \ \mathbb{1}_{(cl_t \in C1)}}{\sum_{t=1}^{n} |ds_t|^p \ \mathbb{1}_{(cl_t \in C1)}}$$

and

$$F_i^{C0} = \frac{\sum_{t=1}^{i} \ \mathbb{1}_{(cl_t \in C0)}}{n - n_a}$$

We set the exponent parameter $p = 1$ so that in the calculation of ES we weight the gene A altered cell lines by their dependence score normalized by the sum of the dependence scores over all of the cell lines in our subset.

We estimate the statistical significance (nominal p-value) of the ES comparing it with the set of scores $ES_{NULL}$ computed with randomly assigned gene A alteration status and reordered cell line list. We perform this permutation step 200 times, recompute the ES of the gene set for the permuted data and compile all results to generate a null distribution for the ES. The empirical, nominal p-value of the observed ES is then calculated relative to this null distribution.

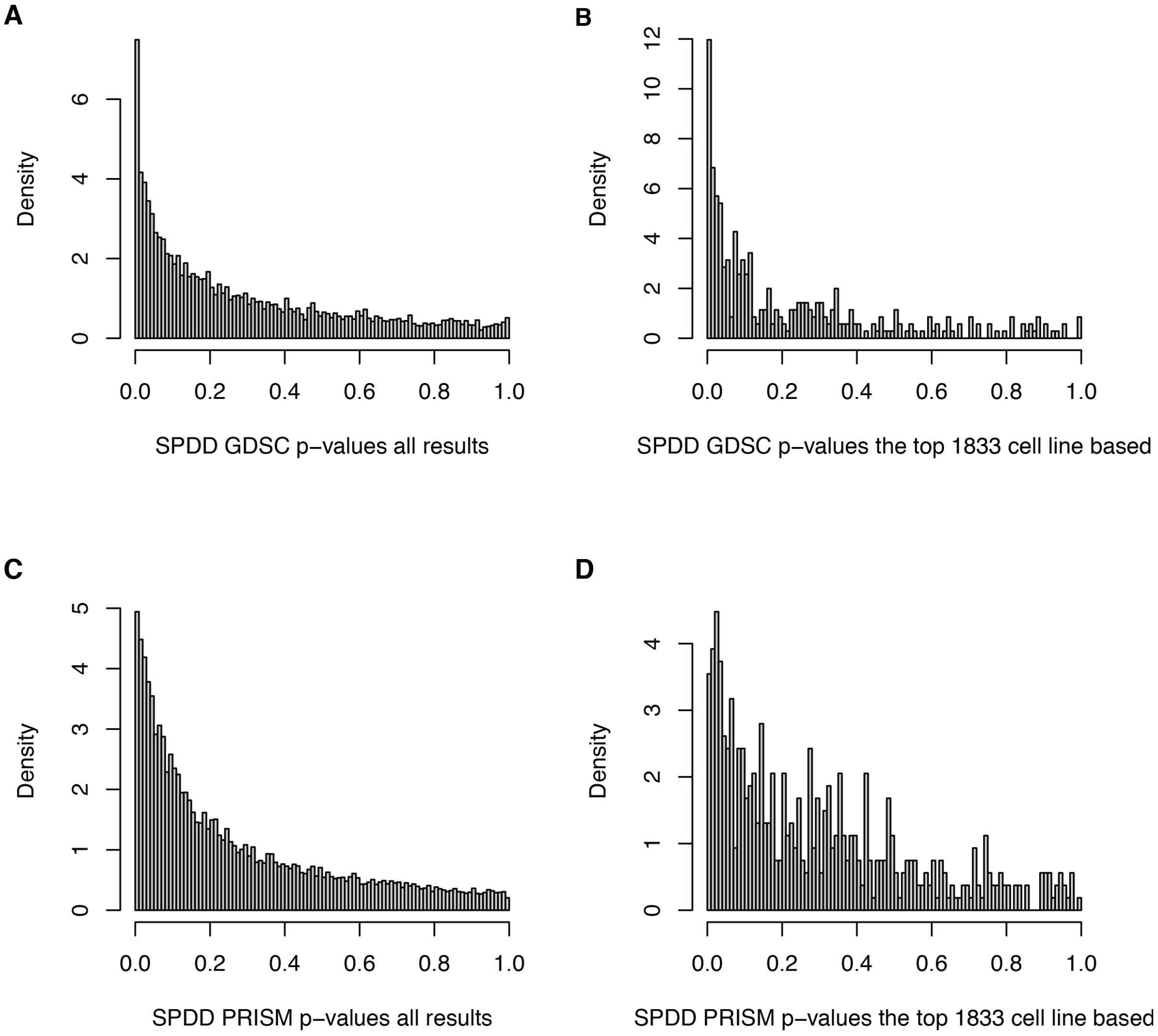


**Supplementary Figure 1: The p-value histograms illustrating the SPDD tests results.** **A** Histogram of all (26,354) p-values of SPDD test done for GDSC dataset **B** Histogram of 351 p-values of SPDD test done for GDSC dataset (among the top 1,833 cell line based pairs, 351 gene pairs passed the test application criteria for GDSC dataset) **C** Histogram of all (62,026) p-values of SPDD test done for PRISM dataset **D** Histogram of 536 p-values of SPDD test done for PRISM dataset (among the top 1,833 cell line based pairs, 536 gene pairs passed the test application criteria for PRISM dataset)
